# Quantifying the relationship between within-host dynamics and transmission for viral diseases of livestock

**DOI:** 10.1101/2023.05.05.539521

**Authors:** Simon Gubbins

**Affiliations:** The Pirbright Institute, Ash Road, Pirbright, Surrey GU24 0NF, U.K

**Keywords:** mathematical modelling, Bayesian methods, foot-and-mouth disease, swine influenza, cattle, pigs

## Abstract

Understanding the population dynamics of an infectious disease requires linking within-host dynamics and between-host transmission in a quantitative manner, but this is seldom done in practice. Here a simple phenomenological model for viral dynamics within a host is linked to between-host transmission by assuming that the probability of transmission is related to log viral titre. Data from transmission experiments for two viral diseases of livestock, foot-and-mouth disease virus in cattle and swine influenza virus in pigs, are used to parameterise the model and, importantly, test the underlying assumptions. The model allows the relationship between within-host parameters and transmission to be determined explicitly through their influence on the individual reproduction number and generation time. Furthermore, these critical within-host parameters (time and level of peak titre, viral growth and clearance rates) can be computed from more complex within-host models, raising the possibility of assessing the impact of within-host processes on between-host transmission in a more detailed quantitative manner.

**Author summary:** For a pathogen to be able to transmit between hosts it must replicate to a sufficiently high level within an infected host. Because of this linking the dynamics of a pathogen within a host to transmission between hosts is important for understanding an infectious disease and its control. In this study I develop a simple mathematical model for the within-host dynamics and combine it with a model relating the probability of transmission to the level of the pathogen. I use the model derive explicit relationships between parameters related to the within-host dynamics, such as viral growth and clearance rates, and summary transmission measures, such as the reproduction number and generation time. I test the assumptions in the underlying model and estimate parameters using data from transmission experiments for two important viral diseases, foot-and-mouth disease virus in cattle and swine influenza virus in pigs. Identifying the critical within host parameters that influence transmission allows the impact of within-host processes on between-host transmission to be investigated in a more detailed quantitative manner.

## Introduction

A pathogen must replicate to a sufficiently high level within an infected host for it to be able to sustain ongoing chains of transmission to new hosts. Consequently, linking within-host dynamics and between-host transmission in a quantitative and predictive manner is important for understanding the population dynamics of an infectious disease, yet it is seldom done in practice [1–4]. Furthermore, most studies which have considered the link between pathogen load and transmission have relied on plausible assumptions rather than empirical data [1,2].

The relationship between pathogen, specifically, viral load and transmission has been measured empirically for viruses such as HIV-1 in humans [5,6], dengue virus in humans and mosquitoes [7], porcine reproductive and respiratory syndrome virus in pigs (PRRSV) [8], foot-and-mouth disease virus (FMDV) in cattle [9], herpes simplex virus-2 in humans [10] and influenza virus in humans [11,12] and pigs [13]. Although these studies quantified the relationship, they did not link it to viral dynamics within the host. More recently, a simple phenomenological model for the within-host dynamics of FMDV was linked to the outcome of environmental transmission experiments [14]. More detailed within-host models for SARS-CoV-2 were linked to transmission using detailed contact data [15,16] or by estimating the contact distribution based on estimated individual reproduction numbers for cases [17]. Finally, a within-host model has been linked with a model for the probability of transmission from mammalian hosts to mosquito vectors for Zika virus [18] and Rift Valley fever virus [19].

These modelling approaches made the link between within-host dynamics and transmission explicit, but they did not explore how processes at one scale (i.e. within-host dynamics) influence those at another (i.e. between-host transmission) in detail (cf. [20,21]). For example, they did not quantify how viral growth or clearance rates impact the individual reproduction number (i.e. the expected number of secondary infections arising from the individual) or the generation time (i.e. the interval between an individual becoming infected and it infecting others).

In this paper, a simple phenomenological model for within-host viral dynamics [14,22,23] was linked to between-host transmission by assuming that the probability of transmission is related to viral titre [2]. This model was then used to derive expressions relating the within-host and transmission parameters to the individual reproduction number and the generation time. The model was parameterised and assumptions tested using data from a series of one-to-one transmission experiments for two viruses that infect livestock: FMDV in cattle [24] and swine influenza virus (SwIV) in pigs [13]. Importantly, these experiments used viruses in their natural hosts and animals that were infected by a natural route (contact with infected animals), rather than inoculation, thereby making them less artificial and more reflective of a real-world situation. Furthermore, the data allowed heterogeneity amongst animals in within-host dynamics and transmission to be explored.

## Results

### Within-host viral dynamics

Within-host viral titres typically rise exponentially following infection, reaching a maximum level after which they decay exponentially. This pattern can be captured using a simple phenomenological model [13,22,23] in which the viral titre at τ days post infection is given by,

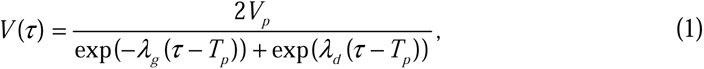

where *V_p_*, *T_p_*, λ*_g_*and λ*_d_* are the peak titre, the time of peak titre and the rates during the exponential growth and decay phases, respectively. Individual variation in within-host viral dynamics can be incorporated by allowing each of the parameters (i.e. *V_p_*, *T_p_*, λ*_g_*and λ*_d_*) to vary amongst individuals. Furthermore, for FMDV the viral dynamics described by equation (1) were linked to the onset of clinical signs by assuming the time of peak titre and incubation period were jointly distributed.

Fitted curves of viral titres over time are shown for FMDV in three compartments (blood, nasal fluid (NF) and oesophageal-pharyngeal fluid (OPF)) in Fig 1 and for SwIV in nasal swabs in Fig 2. The estimates for the within-host viral parameters for FMDV and SwIV are presented in Fig 3. The dynamics of FMDV in blood and NF differed amongst cattle in both peak titre and timing of peak titre, while those in OPF were more consistent in the timing of peak titre (Fig 1). This was reflected in differences amongst animals in all four within-host parameters for blood and NF, whereas only the peak titre in OPF varied greatly amongst individuals (Fig 3). Similarly, the principal difference amongst pigs in within-host parameters for SwIV was in peak titre, while the remaining parameters were consistent amongst individuals (Figs 2 & 3).

**Figure 1.**
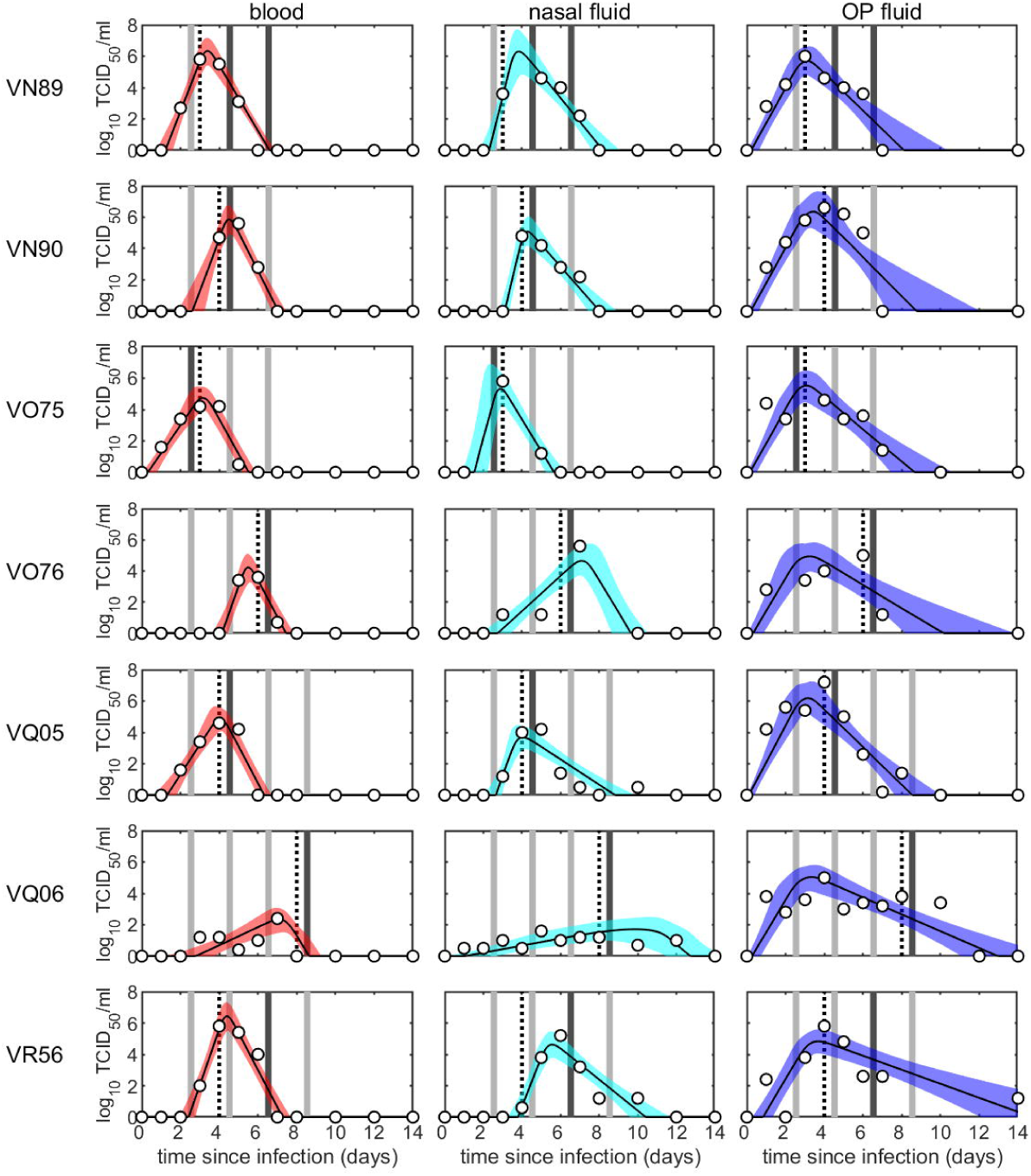
Within-host viral dynamics and transmission of foot-and-mouth disease virus in cattle. Each plot shows the viral titre (log_10_ tissue culture (TC) ID_50_/ml) in blood (red), nasal fluid (cyan) or oesophageal-pharyngeal (OP) fluid (blue) for the animal (identified to the left of the first column): posterior median (solid black line) and 2.5th and 97.5th percentiles (shading) for the fitted virus curves, (1). Timing and outcome of experimental challenges are indicated by bars, which are coloured dark grey if the challenge was successful and light grey if it was not. Open circles give the observed viral titres for each animal. The vertical black dashed lines indicate the onset of clinical signs.

**Figure 2.**
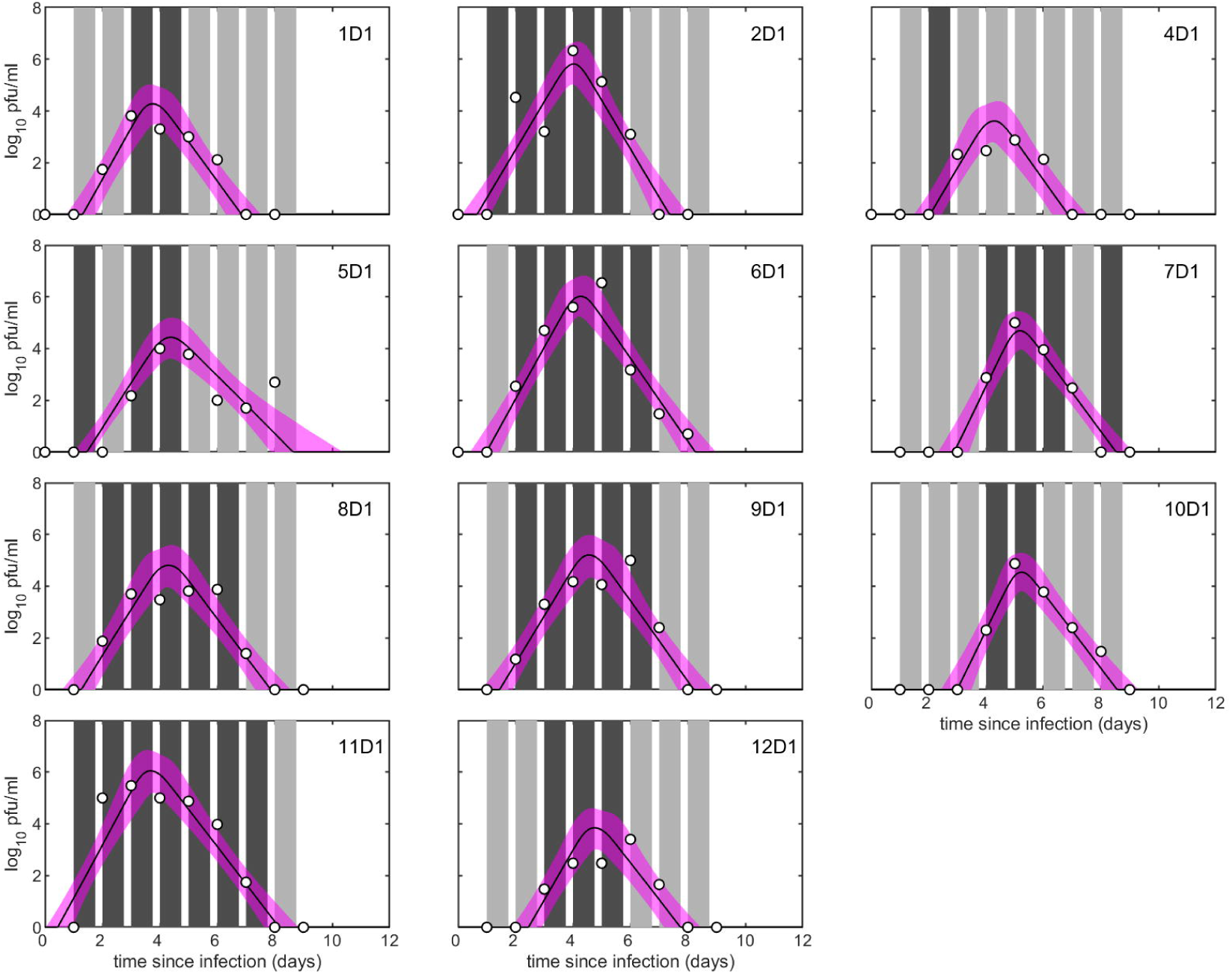
Within-host viral dynamics and transmission of swine influenza virus in pigs. Each plot shows the viral titre (log_10_ pfu/ml) in nasal swabs for the animal (identified in the top right-hand corner): posterior median (solid black line) and 2.5th and 97.5th percentiles (shading) for the fitted virus curves, (1). Timing and outcome of experimental challenges are indicated by bars, which are coloured dark grey if the challenge was successful and light grey if it was not. Open circles give the observed viral titres for each animal.

**Figure 3.**
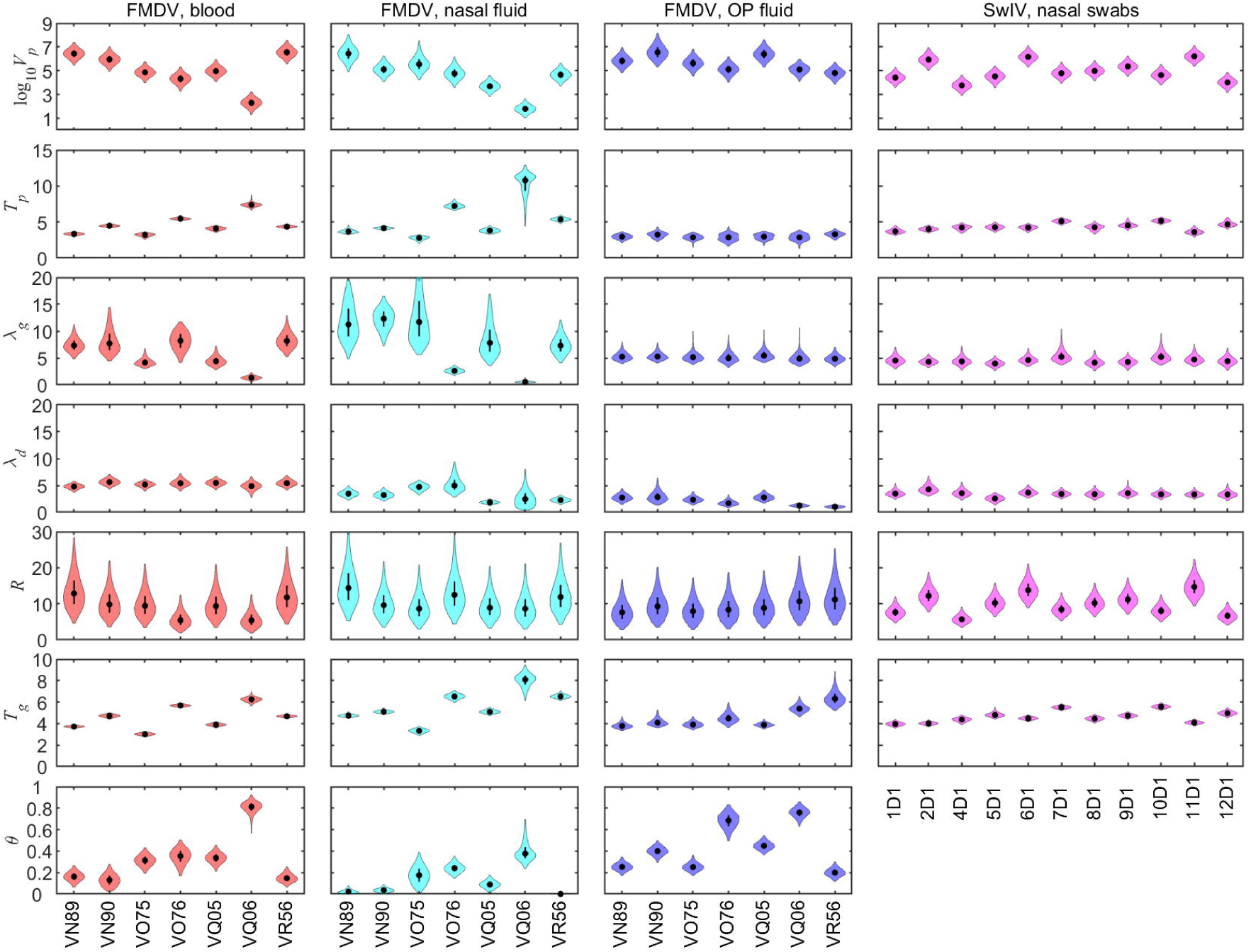
Within-host parameters and summary transmission measures for cattle infected with foot-and-mouth disease virus (FMDV) and pigs infected with swine influenza virus (SwIV). Within-host parameters are: peak titre (log_10_ *V_p_*; log_10_ tissue culture ID_50_/ml for cattle, log_10_ pfu/ml for pigs); time of peak titre (*T_p_*; days post infection); rate for the exponential viral growth phase (λ*_g_*; per day); and rate for the exponential viral decay phase (λ*_d_*; per day). Summary transmission measures are: individual reproduction number (*R*); generation time (*T_g_*; days); and proportion of transmission before the onset of clinical signs (θ). Violin plots show the posterior density (shape), the posterior median (black circles) and the interquartile range (black line). Shape colour indicates the virus and the viral titre used in the model: FMDV and viral titre in blood (red), nasal fluid (cyan) or oesophageal-pharyngeal (OP) fluid (blue); or SwIV and viral titre in nasal swabs (magenta). Because none of the SwIV-infected pigs showed clinical signs, θ was not calculated in this case.

### Linking within-host dynamics and transmission

Within-host viral dynamics can be linked to between-host transmission by assuming that the probability of transmission is related to viral titre. Assuming frequency dependent- transmission, which is appropriate for farm animals [25,26], the probability of transmission from an infected to a susceptible animal when exposure occurs between τ_0_ and τ_1_ days post infection is given by,

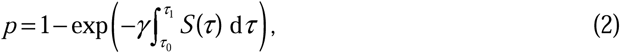

where γ is the transmission parameter and *S*(τ) is the level of viral shedding at τ days post infection.

Viral shedding was assumed to be proportional to log titre (i.e. *S*(τ)=log *V*(τ), where *V*(τ) is given by equation (1)) and the transmission parameter (γ) was assumed to be the same for all animals. Alternative models were considered in which shedding was assumed to be proportional to titre or in which the transmission parameter varied amongst animals. However, none of these was better supported by the data (S1 Table). Model fits and parameter estimates for the alternative models are discussed in S1 Text and compared in S1- S3 Figs.

Virus isolation from NF was the best proxy measure for FMDV infectiousness. The model using NF as the proxy adequately captured the challenge outcomes for all seven animals (posterior predictive *P*-values >0.15 for all animals) and had the highest posterior predictive *P*-value in a majority (5 out of 7) of comparisons (S2 Table). By contrast, there were animals for which the models using virus isolation from blood or from OPF as the proxy were not reliably able to capture the challenge outcomes (i.e. posterior predictive *P*-values <0.05) (S2 Table; cf. Fig 1).

### Relationship between within-host dynamics and transmission

To explore the relationship between within-host viral dynamics and between-host transmission two measures summarising transmission between individuals were considered: the individual reproduction number (*R*); and the generation time (*T_g_*). Between-animal variation in the within-host parameters resulted in between-animal variation in the individual reproduction numbers and generation times (Figs 3 & 4; see also S4-S7 Figs). In addition, for FMDV the individual reproduction number for an animal was broadly similar for each of the proxy measures of infectiousness (Fig 3; S4 Fig). By contrast, the generation time for an animal was consistently higher when using NF as a proxy compared with blood, but with no clear pattern for OPF (Fig 3; S6 Fig).

**Figure 4.**
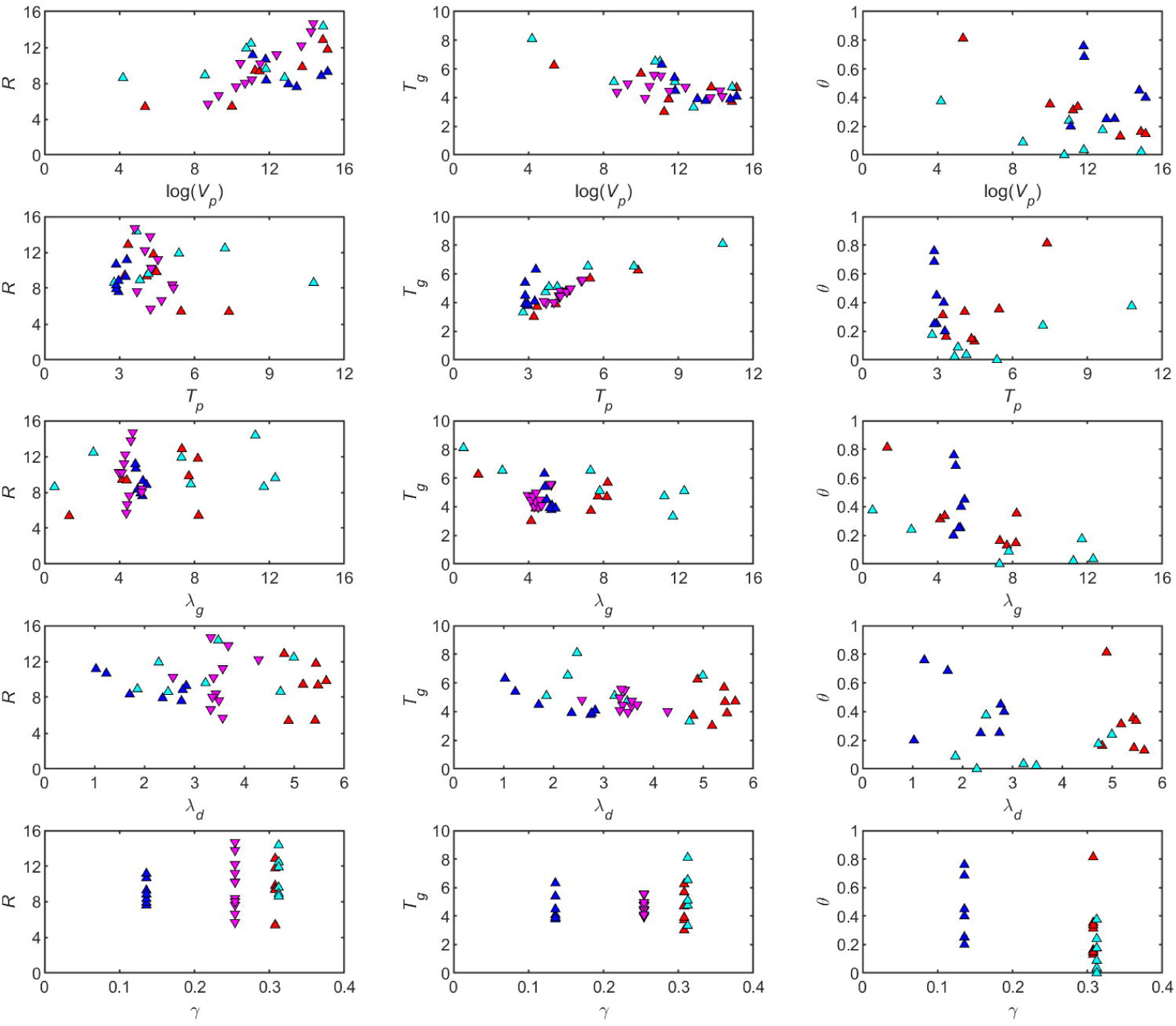
Summary transmission measures for foot-and-mouth disease virus (FMDV) in cattle and swine influenza virus (SwIV) in pigs and their dependence on within-host viral dynamics. Plots show the individual reproduction number (*R*; left-hand column), generation time (*T_g_*; middle column) or proportion of transmission before the onset of clinical signs (θ; right-hand column) and its dependence on peak titre (log *V_p_*), the time of peak titre (*T_p_*) and the rates during the exponential viral growth (λ*_g_*) and decay (λ*_d_*) phases and the transmission parameter (γ). Shape and colour indicate the virus and the viral titre used in the model: FMDV and viral titre in blood (red up-triangles), nasal fluid (cyan up-triangles) or oesophageal-pharyngeal (OP) fluid (blue up-triangles); or SwIV and viral titre in nasal swabs (magenta down-triangles). Because none of the SwIV-infected pigs showed clinical signs, θ was not calculated in this case.

Based on a visual inspection there were no clearly discernible relationships between most within-host parameters and either the individual reproduction number or generation time (Fig 4). However, using a heuristic approximation to the virus shedding curve expressions were derived relating the individual reproduction number, *R*, and generation time, *T_g_*, to the within-host parameters (*V_p_*, *T_p_*, λ*_g_* and λ*_d_*) and transmission parameter (γ) (S2 Text). For the individual reproduction number, this relationship is

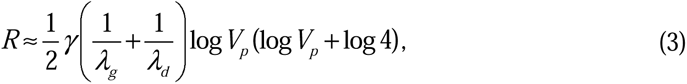

indicating that *R* increases with an increase in transmission parameter (γ) or peak viral titre (*V_p_*) and decreases with an increase in rates in the exponential growth or decay phases (λ*_g_* or λ*_d_*), but is independent of the time of peak viraemia (*T_p_*). The corresponding relationship between the generation time and the within-host and transmission parameters is

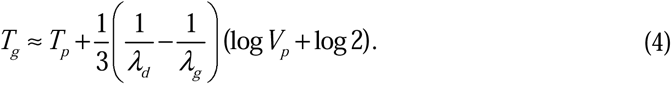

This shows that *T_g_* increases as the time and level of peak titre (*T_p_*and *V_p_*) and the rate in the exponential growth phase (λ*_g_*) increase and decreases as the rate in the exponential decay phase (λ*_d_*) increases, but is independent of the transmission parameter (γ).

The relationship with *R* or *T_g_*inferred heuristically is apparent when some of the within-host parameters are considered in isolation (e.g. *R* and log *V_p_* or *T_g_* and *T_p_*), but less so for others because of the dependence on the remaining parameters (Fig 4). However, the relationships are clearer when considering all sampled parameters (S4-S7 Figs) rather than just their median values (Fig 4). In addition, the direction of the relationship (i.e. whether an increase in the parameter is associated with an increase or decrease in *R* or *T_g_*) is supported by the partial rank correlation coefficients between the parameters and *R* (S4 & S5 Figs) or *T_g_* (S6 & S7 Figs).

### Effectiveness of reactive control measures

For FMDV, the proportion of transmission before the onset of clinical signs (denoted by θ) was also calculated for each animal, which is useful for assessing the efficacy of reactive control measures [24,27]. Again, between-animal variation in within-host dynamics resulted in between-animal variation in θ (Figs 3 & 4). Based on partial rank correlation coefficients, larger values of θ are associated with earlier times of peak titre and lower rates in the exponential growth phase (S8 Fig). Furthermore, for each animal the smallest value of θ was obtained when NF was used as the proxy measures of infectiousness, followed by blood and then OPF (Fig 3; S8 Fig).

## Discussion

In this study the relationship between within-host dynamics and between-host transmission was quantified for two viral diseases of livestock, FMDV in cattle and SwIV in pigs, using empirical data from transmission experiments, thereby allowing underlying assumptions in the modelling approach to be tested. Explicit, if approximate, relationships were derived between the within-host (*T_p_*, log *V_p_*, λ*_g_*, λ*_d_*) and transmission (γ) parameters and the individual reproduction number and generation time (*R* and *T_g_*, given by equations (3) and (4), respectively).

The four parameters in the model used to describe the within-host dynamics (*T_p_*, log *V_p_*, λ*_g_*, λ*_d_*) each reflect the net effect of a combination of underlying biological processes and so mask some of the inherent complexity of the within-host dynamics. For example, the level and timing of peak viral titre is likely to be related to the innate immune response, particular the interferon response, while viral clearance relates to the levels of antibody and T cell responses (see [24,28] for FMDV and [29,30] for SwIV). This suggests that *T_p_*, log *V_p_* and λ*_g_* are likely to be influenced primarily by innate immune responses, while λ*_d_*is likely to be influenced primarily by adaptive immune responses.

The choice of modelling approach reflects in part a limitation of using only viral titres when estimating within-host parameters, such that only the four parameters in the model are identifiable from the data [22,31]. The model is nonetheless able to capture the trajectories of the viral titres within a host (Figs 1 & 2) and a similar phenomenological approach has been used previously for FMDV [14], influenza virus [22,23] and SARS-CoV-2 [32,33].

More complex models have been developed to describe the within-host dynamics of viruses, including influenza virus [29,34] and FMDV [28,35]. These models incorporate features of the within-host biology, including viral replication in populations of cells and both innate, especially the interferon response, and adaptive immune responses. Parameters in these more complex models can be related to those in the phenomenological virus curve given by equation (1) using approximation methods [36,37]. This provides a means of simplifying a more complex within-host model so that it can be embedded in a between-host transmission model [1]. Furthermore, it makes it easier to assess the impact of within-host processes (e.g. viral replication, innate or adaptive immune responses) on between-host transmission in a quantitative manner.

Transmission does not depend solely on the within-host dynamics of a pathogen. Rather it also depends on additional factors, including host behaviour and frequency and duration of contacts, the effects of which are incorporated implicitly in the transmission parameter [38]. The best supported models for both FMDV and SwIV assumed a common transmission parameter amongst animals (S1 Table), implying that between-host variation in these factors is of limited importance in transmission experiments, especially the one-to-one design used in the studies analysed here. This is unlikely to be the case in the field, however, where contacts between animals are likely to be more variable [39–41].

Although there was no evidence for contact heterogeneity in the transmission experiments, there was still substantial variation amongst animals in their summary transmission measures (Fig 4). This reflects variation amongst the animals in within-host dynamics for both FMDV and SwIV (Fig 3). The level of variation in the parameters differed amongst viruses and compartments, with the most consistently variable parameter being peak viral titre. Despite this variation, however, all animals had individual reproduction numbers well above one (Fig 4), suggesting that variation at the within-host level may not result in much variation at the population level in this case. This is in contrast with other viruses for which there can be substantial differences amongst individuals in terms of infectiousness [42]. Such variation can arise through heterogeneities in viral dynamics, as well as in contacts [16,17].

When scaling from within-host dynamics to between-host transmission there are two important issues to consider [1,2]. First, how does viral load relate to shedding and, hence, transmission? Second, what is the most appropriate proxy measure (i.e. viral titre in which compartment) for infectiousness? Model selection suggested that, for FMDV in cattle and SwIV in pigs, models in which viral shedding is proportional to log titre were better supported by the data than ones in which shedding is proportional to titre, particularly if the transmission parameter was common to all animals (S1 Table). A similar conclusion was also reached for SARS-CoV-2 [15]. Assumptions about shedding have a large impact on estimates for the individual reproduction number, which can be several orders of magnitude higher (and sometimes unrealistically high) when shedding is proportional to titre compared with when it is proportional to log titre (S3 Fig).

The analyses for FMDV show that the choice of proxy measure used for infectiousness (i.e. viral titres in different compartments) did not influence model selection (S1 Table) and, hence, inferences about how transmission relates to viral titre and variation between animals in transmission parameter. However, the choice of proxy measure did influence the estimates of the summary transmission measures and the predictions of the effectiveness of reactive control measures (Figs 3 & 4; S4, S6 & S8 Figs). Consequently, it is essential to determine which, if any, of the proxy measures considered are reliable indicators of infectiousness.

## Conclusion

In this study, explicit expressions have been derived that relate viral dynamics within a host (level and time of peak titre, virus growth and clearance rates) to between-host transmission (individual reproduction number and generation time). These expressions have been parameterised and the underlying assumptions tested using empirical data from transmission experiments for two viral diseases of livestock, FMDV in cattle and SwIV in pigs. The modelling approach provides a means of embedding more complex within-host models in between-host transmission models. Furthermore, the critical within-host parameters (time and level of peak titre, viral growth and clearance rates) can be computed from more complex within-host models. This will facilitate assessing the impact of within-host processes on between-host transmission in a more detailed quantitative manner, including the impact of intervention strategies, for example, the use of anti-virals.

## Methods

### Data

Parameters in the model given by equations (1) and (2) were estimated using previously-published data from a series of one-to-one transmission experiments for FMDV (O UKG 34/2001) in cattle [24] and SwIV (H1N1pdm09) in pigs [13]. In both cases uninfected recipient animals were challenged by exposure to infected donor animals at multiple times post infection of the donor and the outcome of challenge recorded (i.e. whether or not transmission occurred) (Figs 1 & 2). The donor animals were infected by direct contact challenge with another infected animal, rather than by inoculation. As well as challenge outcome, viral titres were measured daily in the donor animals. In the FMDV experiments titres were measured in three compartments: blood, nasal fluid (NF) and oesophageal- pharyngeal fluid (OPF) (Fig 1). In the SwIV experiments titres were measured daily in nasal swabs (Fig 2). All data are available from the original publications [13,24], but are provided in the supporting information (S1 & S2 Data) in the format used in the present analysis.

### Individual variation in within-host and transmission parameters

Individual variation in within-host viral dynamics, (1), was incorporated by allowing each of the parameters (i.e. *V_p_*, *T_p_*, λ*_g_* and λ*_d_*) to differ amongst individuals. Specifically, the parameters were assumed to be drawn from higher-order distributions, so that,

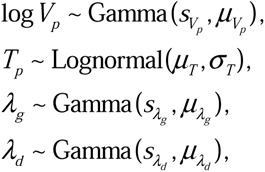

where *s_i_* and μ*_i_* (*i*=*V_p_*,λ*_g_*,λ*_d_*) are the shape parameter and mean of the gamma distributions, respectively, and μ*_T_* and σ*_T_* are the parameters for the log normal distribution (mean and standard deviation on the log scale, respectively). Estimates for the hierarchical parameters are presented in S3 Table for FMDV and in S4 Table for SwIV.

In addition, the viral dynamics described by equation (1) were linked to the onset of clinical disease by assuming the time of peak titre and incubation period follow a bivariate log normal distribution. In this case, the time of onset of clinical signs, *C*, can be written conditionally on the time of peak titre, so that,

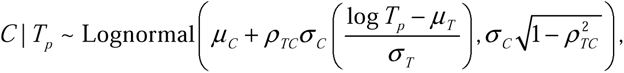

where μ*_i_* and σ*_i_* (*i*=*C*,*T*) are the parameters for the (marginal) log normal distributions (mean and standard deviation on the log scale, respectively) and ρ*_TC_* is the correlation between the times of peak titre and clinical onset. None of the pigs infected with SwIV showed clinical signs [13], hence, parameters related to clinical disease (μ*_C_*, σ*_C_*, ρ*_TC_* and θ) were estimated only for FMDV (S3 Table).

When the transmission parameter varied amongst individuals, the parameter for each animal was drawn from a higher-order distribution. When shedding was proportional to titre, the log-transformed transmission parameters were drawn from a normal distribution (i.e. log γ*_j_*∼Normal(μ_γ_,σ_γ_), where μ_γ_ and σ_γ_ are the mean and standard deviation, respectively). When shedding was proportional to log titre, the transmission parameters were drawn from a gamma distribution (i.e. γ*_j_*∼Gamma(*s*_γ_,μ_γ_), where *s*_γ_ and μ_γ_ are the shape parameter and mean, respectively).

### Parameter estimation

Parameters were estimated using Bayesian methods. The likelihood for the data (comprising the challenge outcomes, δ*_ij_*, the virus isolation data, *V*_j_ ^(*obs*)^(*t*_j_), and, for FMDV, the times of clinical onset, *C_j_* for donor animal *j*) is given by,

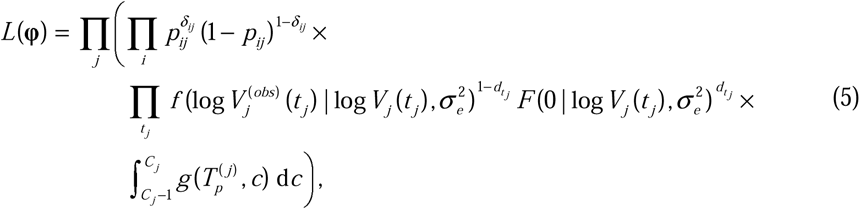

where ϕ is a vector of model parameters, *p_ij_* is the probability of transmission at the *i*th challenge for donor animal *j* (given by equation (2)), *V_j_* is the viral titre at time *t_j_* (given by equation (1)), *f* and *F* are the probability and cumulative density functions for the normal distribution, respectively, *d* is a variable indicating whether (*d*=1) or not (*d*=0) the observation is left-censored (i.e. it is below the detection threshold, set arbitrarily at 1 TCID_50_/ml (FMDV) or 1 pfu/ml (SwIV)) and *g* is the probability density function for the bivariate log normal distribution. The prior distributions used for each parameter are presented in S5 Table.

Samples from the joint posterior density were generated using an adaptive Metropolis scheme [43], modified so that the scaling factor was tuned during burn-in to ensure an acceptance rate of between 20% and 40% for more efficient sampling of the target distribution [44]. Two chains of 10,000,000 iterations were run, with the first 5,000,000 iterations discarded to allow burn-in of the chains. Each chain was subsequently thinned by taking every 500th iteration.

The adaptive Metropolis scheme was implemented in Matlab (version R2020b; The Mathworks Inc.) and the code is available online [45]. Convergence of the scheme was assessed visually and by examining various criteria provided in the coda package [46] in R (version 4.0.5) [47].

The four models for viral shedding (i.e. proportional to titre or log titre) and variation in transmission parameters amongst animals (i.e. common to all animals or varies amongst animals) were compared using the deviance information criterion (DIC) [48].

For FMDV, the models using viral titres in different compartments (i.e. blood, NF or OPF) were compared by computing the posterior predictive *P*-values for transmission by each animal. Specifically, the joint posterior distribution for an animal was sampled and the probability of transmission at each challenge computed. Whether or not transmission occurred at each challenge was then simulated and the observed outcomes of all challenges for the animal were compared to the simulated ones. This was repeated multiple times and the proportion of samples for which the observed and simulated outcomes matched was computed (i.e. the posterior predictive *P*-value).

### Summary transmission measures

Assuming frequency dependent-transmission, which has been shown to be appropriate for farm animals [25,26], the individual reproduction number, *R*, the generation time, *T_g_*, and the proportion of transmission before the onset of clinical signs, θ, are given by,

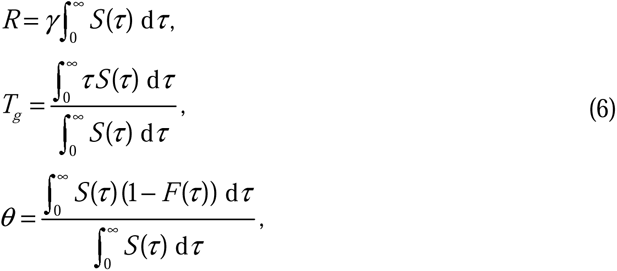

where γ is the transmission parameter, *S*(τ) is the level of virus shedding by an individual at time τ post infection and *F*(τ) is the cumulative density function for the incubation period conditional on the time of peak titre.

The relationship between these measures and the within-host parameters (*V_p_*, *T_p_*, λ*_g_* and λ*_d_*) and the transmission parameter (γ) were explored by computing the values of *R*, *T_g_* and θ for each sample from the joint posterior distribution using the expressions in equation (6). Furthermore, a heuristic approximation to the best-supported shedding curve, *S*(τ)=log *V*(τ) (S9 Fig), was used to derive the explicit relationships between *R* or *T_g_*and *V_p_*, *T_p_*, λ*_g_*, λ*_d_*and γ given by equations (3) and (4) (see S2 Text for details). The accuracy of these approximations was assessed by comparing them to the exact values computed using the expressions in equation (6) (S6 Table).

The sensitivity of the summary transmission measures (*R*, *T_g_* and θ) to changes in the within-host and transmission parameters was further explored by computing partial rank correlation coefficients [49] using samples drawn from the joint posterior distribution for the parameters.

## Supporting information

S1 Text: Additional results

S2 Text: Mathematical derivation of R0 and Tg

S1 Table: Model selection results

S2 Table: Posterior predictive P-values

S3 Table: Parameter estimates (FMDV)

S4 Table: Parameter estimates (SwIV)

S5 Table: Prior distributions

S6 Table: Approximation accuracy

## Ethical information

No animal experiments were carried out as part of the present study. The data used in the analyses come from animal experiments that were conducted in accordance with the UK Animals (Scientific Procedures) Act (ASPA) 1986 and associated guidelines and were approved by the Animal Welfare and Ethical Review Board of The Pirbright Institute.

## Data availability

The data-sets used in the study are available from the original publications and are also provided in the supplementary material. All code used in the analyses is available online [45].

## Author contributions

Conceptualization: SG. Formal analysis: SG. Methodology: SG. Software: SG. Writing - original draft: SG. Writing - review & editing: SG.

## Acknowledgements

The author is grateful to David Schley (formerly at The Pirbright Institute) for helpful discussions at earlier stages in the development of this work.

## Supporting information

**S1 Data. Data on virus isolation and challenge outcome from a series of one-to-one transmission experiments for foot-and-mouth disease virus in cattle.**

**S2 Data. Data on virus isolation and challenge outcome from a series of one-to-one transmission experiments for swine influenza virus in pigs.**

**S1 Fig. Within-host viral dynamics of foot-and-mouth disease virus in cattle.** Circles show the observed viral titre (log_10_ tissue culture (TC) ID_50_/ml) in blood (red), nasal fluid (cyan) or oesophageal-pharyngeal (OP) fluid (blue) for each animal (identified to the left of the first column). Lines show the posterior median for the fitted virus curves, (1), for four models assuming: shedding proportional to titre, transmission parameter common to all animals (black dotted lines); shedding proportional to titre, transmission parameter varies amongst animals (black dashed line); shedding proportional to log titre, transmission parameter common to all animals (black solid line); and shedding proportional to log titre, transmission parameter varies amongst animals (black dash-dotted line).

**S2 Fig. Within-host viral dynamics of swine influenza virus in pigs.** Circles show the observed viral titre (log_10_ pfu/ml) in nasal swabs for each animal (identified in the top right-hand corner). Lines show the posterior median for the fitted virus curves, (1), for four models assuming: shedding proportional to titre, transmission parameter common to all animals (black dotted lines); shedding proportional to titre, transmission parameter varies amongst animals (black dashed line); shedding proportional to log titre, transmission parameter common to all animals (black solid line); and shedding proportional to log titre, transmission parameter varies amongst animals (black dash-dotted line).

**S3 Fig. Within-host parameters, transmission parameters and summary transmission measures for cattle infected with foot-and-mouth disease virus (FMDV) and pigs infected with swine influenza virus (SwIV)**: peak titre (log_10_ *V_p_*; log_10_ tissue culture ID_50_/ml for cattle, log_10_ pfu/ml for pigs); time of peak titre (*T_p_*; days post infection); rate for the exponential viral growth phase (λ*_g_*; per day); rate for the exponential viral decay phase (λ*_d_*; per day); transmission parameter (γ); individual reproduction number (*R*); generation time (*T_g_*; days); and proportion of transmission before the onset of clinical signs (θ). Plots show the posterior median (symbols) and 2.5th and 97.5th percentiles (black line). Symbols indicate model assumptions: shedding proportional to titre, transmission parameter common to all animals (circles); shedding proportional to titre, transmission parameter varies amongst animals (down-triangles); shedding proportional to log titre, transmission parameter common to all animals (diamonds); and shedding proportional to log titre, transmission parameter varies amongst animals (up-triangles). Because none of the SwIV-infected pigs showed clinical signs, θ was not calculated in this case.

**S4 Fig. Individual reproduction number (*R*) and its dependence on within-host viral dynamics for seven cattle infected with foot-and-mouth disease virus.** The first five columns show *R* and its dependence on peak titre (log *V_p_*), the time of peak titre (*T_p_*) and the rates during the exponential viral growth (λ*_g_*) and decay (λ*_d_*) phases and on the transmission parameter (γ). Coloured dots show the parameters for each animal (indicated to the left of the first column) sampled from the joint posterior distribution when using viral titre in blood (red), nasal fluid (cyan) or oesophageal-pharyngeal fluid (blue) to estimate shedding. White crosses mark the posterior median and white lines show the relationship between *R* and the parameter as defined by equation (3), with the remaining parameters fixed at their posterior median values. The sixth (right-hand) column shows partial rank correlation coefficients (PRCC) between the within-host and transmission parameters and *R* calculated for each compartment.

**S5 Fig. Individual reproduction number (*R*) and its dependence on within-host viral dynamics for eleven pigs infected with swine influenza virus.** The first five columns show *R* and its dependence on peak titre (log *V_p_*), the time of peak titre (*T_p_*) and the rates during the exponential viral growth (λ*_g_*) and decay (λ*_d_*) phases and on the transmission parameter (γ). Coloured dots show the parameters for each animal (indicated to the left of the first column) sampled from the joint posterior distribution. White crosses mark the posterior median and white lines show the relationship between *R* and the parameter as defined by equation (3), with the remaining parameters fixed at their posterior median values. The sixth (right-hand) column shows partial rank correlation coefficients (PRCC) between the within-host and transmission parameters and *R*.

**S6 Fig. Generation time (*T_g_*) and its dependence on within-host viral dynamics for seven cattle infected with foot-and-mouth disease virus.** The first five columns show *T_g_* and its dependence on peak titre (log *V_p_*), the time of peak titre (*T_p_*) and the rates during the exponential viral growth (λ*_g_*) and decay (λ*_d_*) phases and on the transmission parameter (γ). Coloured dots show the parameters for each animal (indicated to the left of the first column) sampled from the joint posterior distribution when using viral titre in blood (red), nasal fluid (cyan) or oesophageal-pharyngeal fluid (blue) to estimate shedding. White crosses mark the posterior median and white lines show the relationship between *T_g_* and the parameter as defined by equation (4), with the remaining parameters fixed at their posterior median values. The sixth (right-hand) column shows partial rank correlation coefficients (PRCC) between the within-host and transmission parameters and *T_g_* calculated for each compartment.

**S7 Fig. Generation time (*T_g_*) and its dependence on within-host viral dynamics for eleven pigs infected with swine influenza virus.** The first five columns show *T_g_*and its dependence on peak titre (log *V_p_*), the time of peak titre (*T_p_*) and the rates during the exponential viral growth (λ*_g_*) and decay (λ*_d_*) phases and on the transmission parameter (γ). Coloured dots show the parameters for each animal (indicated to the left of the first column) sampled from the joint posterior distribution. White crosses mark the posterior median and white lines show the relationship between *T_g_* and the parameter as defined by equation (4), with the remaining parameters fixed at their posterior median values. The sixth (right-hand) column shows partial rank correlation coefficients (PRCC) between the within-host and transmission parameters and *T_g_*.

**S8 Fig. Proportion of transmission before the onset of clinical signs (**θ**) and its dependence on within-host viral dynamics for seven cattle infected with foot-and-mouth disease virus.** The first five columns show θ and its dependence on peak titre (log *V_p_*), the time of peak titre (*T_p_*) and the rates during the exponential viral growth (λ*_g_*) and decay (λ*_d_*) phases and on the transmission parameter (γ). Coloured dots show the parameters for each animal (indicated to the left of the first column) sampled from the joint posterior distribution when using viral titre in blood (red), nasal fluid (cyan) or oesophageal-pharyngeal fluid (blue) to estimate shedding. White crosses mark the posterior medians. The sixth (right-hand) column shows partial rank correlation coefficients (PRCC) between the within-host and transmission parameters and θ calculated for each compartment.

**S9 Fig. Approximating the virus shedding curve.** The plot shows the virus shedding curve (blue dashed line) and its piecewise linear approximation (solid red line) (see S2 Text for details). The red shaded area indicates the approximation to the area under the blue dashed shedding curve.

**S1 Table. Deviance information criterion comparing different models for viral shedding and transmission parameters for foot-and-mouth disease virus (FMDV) in cattle and swine influenza virus (SwIV) in pigs.**

**S2 Table. Posterior predictive *P*-values assessing the titre of foot-and-mouth disease virus in different compartments as a proxy for infectiousness.**

**S3 Table. Summary statistics for the marginal posterior densities for hierarchical parameters in a model linking within-host dynamics and transmission of foot-and-mouth disease virus in cattle.**

**S4 Table. Summary statistics for the marginal posterior densities for hierarchical parameters in a model linking within-host dynamics and transmission of swine influenza virus in pigs.**

**S5 Table. Prior distributions used when estimating parameters related to the within-host dynamics and transmission of foot-and-mouth disease virus (FMDV) in cattle and swine influenza virus (SwIV) in pigs.**

**S6 Table. Accuracy of heuristic approximations for the individual reproduction number and generation time.**

**S1 Text. Alternative models for viral shedding and transmission parameters.**

**S2 Text. Relating summary transmission measures and within-host parameters.**

