## Supplementary material for "Quantifying the relationship between within-host dynamics and transmission for viral diseases of livestock": S1 Text: Additional results

**S1 Text. Alternative models for viral shedding and transmission parameters**

Four models were considered when linking within-host viral dynamics and transmission. These differed in whether viral shedding was assumed to be proportional to titre or log titre and whether the transmission parameter was common to all animals or varied amongst animals. The best-supported model (as judged by the deviance information criterion) for both foot-and-mouth disease virus and swine influenza virus was one in which shedding was proportional to log titre and the transmission rate was common to all animals (S1 Table). Accordingly, this model is the focus of the results presented in the main paper. Here the model fits and parameter estimates for the alternative models are discussed briefly.

*Within-host viral dynamics*. Fitted curves of viral titres over time are shown for FMDV in three compartments (blood, nasal fluid (NF) and oesophageal-pharyngeal fluid (OPF)) in S1 Fig and for swine influenza in nasal swabs in S2 Fig. These are broadly similar across the four models in all cases. The only systematic differences are in the fitted titres for FMDV in OPF, which increase more slowly and peak later for the models in which shedding is assumed to be proportional to titre. This is also reflected in the estimates for the within-host parameters which are similar for all four models (S3 Fig). The exceptions are the estimates for the time of peak titre (*T_p_*) and rate for the exponential growth phase (*λ_g_*) for FMDV in OPF, where *T_p_* is higher and is *λ_g_* lower for the models in which shedding is assumed to be proportional to titre.

*Transmission parameters*. The transmission parameters are much lower when shedding is assumed to be proportional to titre compared with when shedding is assumed to be proportional to log titre (S3 Fig). This reflects the marked difference in the magnitude of the assumed level of shedding in the models. When shedding is assumed to be proportional to log titre, transmission parameters do not differ greatly amongst animals (S3 Fig). By contrast, there is substantial variation in transmission parameters when shedding is assumed to be proportional to titre, except for FMDV where viral titre in OPF is used as the proxy for infectiousness (S3 Fig).

*Summary transmission measures*. The individual reproduction number (*R*) was sensitive to the assumptions made about viral shedding and whether the transmission parameter varied amongst animals (S3 Fig). Estimates for *R* were typically, but not always, higher when shedding was assumed to be proportional to titre. Indeed, some estimates for *R* for this model are unrealistically high (*R*>100). The proportion of transmission before the onset of clinical signs (*θ*) was also sensitive to assumptions about viral shedding, but not about the transmission parameter (S3 Fig). The estimates for *θ* were generally, but not always, higher when shedding was assumed to be proportional to log titre compared with when shedding was assumed to be proportional to titre. By contrast, the generation times (*T_g_*) for each animal were consistent for all four models (S3 Fig).

*Proxy measures of infectiousness for FMDV*. For all four models, virus isolation from NF was the best proxy measure for infectiousness. The model using NF as the proxy adequately captured the challenge outcomes for all seven animals (posterior predictive *P*-values >0.05 for all but one case and >0.15 for all but three cases) and had the highest posterior predictive *P*-value in a majority (21 out of 28) of cases (S2 Table). By contrast, there were animals for which the models using virus isolation from blood or from OPF as the proxy were not reliably able to capture the challenge outcomes (i.e. posterior predictive *P*-values <0.05) (S2 Table).
