## Supplementary material for "Quantifying the relationship between within-host dynamics and transmission for viral diseases of livestock": S2 Text: Mathematical derivation of R0 and Tg

**S2 Text. Relating summary transmission measures and within-host parameters**

Using a heuristic approximation to the virus shedding curve it is possible to derive explicit relationships between the summary transmission measures (*R* and *T_g_*) and the within-host parameters (*V_p_*, *T_p_*, *λ_g_* and *λ_d_*).

***S2.1	Approximating the virus shedding curve***

In the simple phenomenological model used to describe the within-host viral dynamics, the titre, *V*, at *τ* days post infection is given by,

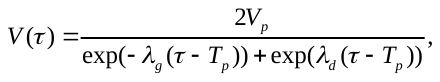

where *V_p_*, *T_p_*, *λ_g_* and *λ_d_* are the peak titre, the time of peak titre and the rates for the exponential viral growth and decay phases, respectively. The corresponding level of viral shedding, *S*(*τ*), is proportional to log titre, so that *S*(*τ*)=log *V*(*τ*) where *V*(*τ*) is the viral titre given by equation . The shedding curve can be approximated by a piecewise linear function (S9 Fig), namely,

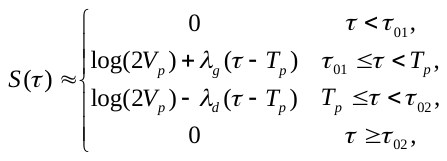

where,

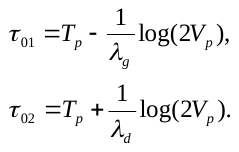

***S2.2	Deriving expressions for the summary transmission measures***

The individual reproduction number, *R*, is given by,

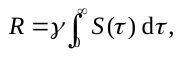

where *γ* is the transmission parameter and *S*(*τ*) is the level of virus shedding. Using equation the integral in equation is approximated by the area of the red trapezium in Figure S9, so that,

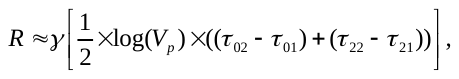

where *τ*_01_ and *τ*_02_ are given by equation and,

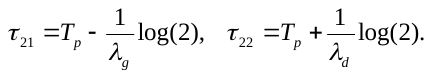

Substituting the expressions for *τ*_01_, *τ*_02_, *τ*_21_ and *τ*_22_ into equation gives the following approximation relating *R* to the within-host parameters,

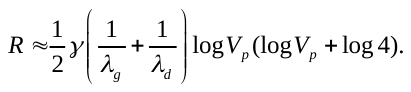

The generation time, *T_g_*, is given by,

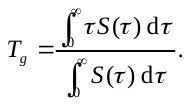

The numerator in equation can be approximated using equation as follows,

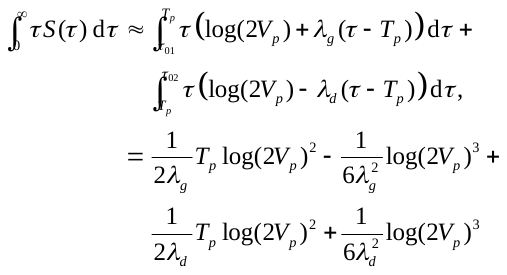

The denominator in equation can be approximated by the large red-bordered triangle in S9 Fig, so that,

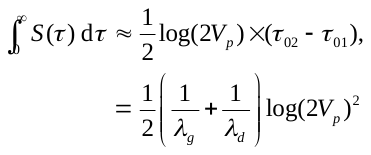

Consequently, the relationship between the within-host parameters and generation time is approximated by,

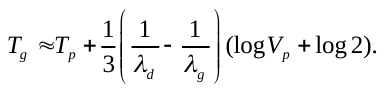

***S2.3	Accuracy of the expressions for the summary transmission measures***

The accuracy of the expressions was checked by comparing the values for the summary transmission measures computed using the explicit expressions, and , for each sample from the joint posterior distribution with those computed using their approximations, and (S6 Table). The approximation slightly overestimates *R* (range of median differences: -0.021 to -0.007) and the absolute differences were all <0.17. The approximation slightly underestimates *T_g_* (range of median differences: 0.001 to 0.005) and the absolute differences were all <0.09. These differences are sufficiently small for it to be reasonable to draw inferences using the expressions for the two measures derived using the approximations.
