## Supplementary material for "Quantifying the relationship between within-host dynamics and transmission for viral diseases of livestock": S1 Table: Model selection results

**S1 Table.** Deviance information criterion (DIC) comparing* different models for viral shedding and transmission parameters for foot-and-mouth disease virus (FMDV) in cattle and swine influenza virus (SwIV) in pigs.

| model |  | FMDV† | | | SwIV |
| --- | --- | --- | --- | --- | --- |
| shedding | transmission | blood | NF | OPF |  |
| proportional to titre | common to all animals | 205.8 | 161.4 | 310.6 | 334.1 |
| proportional to titre | varies amongst animals | 189.3 | 154.3 | 312.0 | 313.6 |
| proportional to log titre | common to all animals | 156.7 | 155.6‡ | 287.7 | 315.3‡ |
| proportional to log titre | varies amongst animals | 160.1 | 159.8 | 291.4 | 314.2 |

* a model with a smaller DIC is preferred to one with a larger DIC

† compartment in which viral titre measured (NF - nasal fluid; OPF - oesophageal-pharyngeal fluid)

‡ because the change in DIC between this model and more complex ones was less than 2, the simpler model was preferred on grounds of model parsimony
