## Supplementary material for "Quantifying the relationship between within-host dynamics and transmission for viral diseases of livestock": S2 Table: Posterior predictive P-values

**S2 Table.** Posterior predictive *P*-values assessing the titre of foot-and-mouth disease virus in different compartments as a proxy for infectiousness.

| model | | compartment‡ | animal ID | | | | | | |
| --- | --- | --- | --- | --- | --- | --- | --- | --- | --- |
| shedding* | transmission† |  | VN89 | VN90 | VO75 | VO76 | VQ05 | VQ06 | VR56 |
| titre | common | blood | 0.000 | 0.992 | 0.539 | 0.041 | 0.754 | 0.003 | 0.002 |
|  |  | NF | 0.065 | 0.944 | 0.867 | 0.822 | 0.413 | 0.011 | 0.603 |
|  |  | OPF | 0.002 | 0.559 | 0.347 | 0.017 | 0.346 | 0.003 | 0.042 |
| titre | varies | blood | 0.000 | 0.955 | 0.663 | 0.322 | 0.743 | 0.011 | 0.001 |
|  |  | NF | 0.311 | 0.906 | 0.833 | 0.847 | 0.665 | 0.078 | 0.676 |
|  |  | OPF | 0.004 | 0.493 | 0.326 | 0.011 | 0.314 | 0.002 | 0.045 |
| log titre | common | blood | 0.009 | 0.504 | 0.344 | 0.278 | 0.333 | 0.031 | 0.059 |
|  |  | NF | 0.251 | 0.392 | 0.356 | 0.365 | 0.303 | 0.167 | 0.212 |
|  |  | OPF | 0.037 | 0.177 | 0.199 | 0.094 | 0.157 | 0.054 | 0.092 |
| log titre | varies | blood | 0.009 | 0.451 | 0.319 | 0.279 | 0.298 | 0.031 | 0.065 |
|  |  | NF | 0.327 | 0.350 | 0.322 | 0.322 | 0.275 | 0.156 | 0.184 |
|  |  | OPF | 0.050 | 0.150 | 0.183 | 0.081 | 0.139 | 0.052 | 0.087 |

* shedding is proportional to viral titre or log viral titre

† transmission parameter is common to all animals or varies amongst animals

‡ compartment in which viral titre measured (NF - nasal fluid; OPF - oesophageal-pharyngeal fluid)
