## Supplementary material for "Quantifying the relationship between within-host dynamics and transmission for viral diseases of livestock": S3 Table: Parameter estimates (FMDV)

**S3 Table.** Summary statistics for the marginal posterior densities for hierarchical parameters in a model linking within-host dynamics and transmission of foot-and-mouth disease virus in cattle.

| parameter | symbol | median | percentiles | |
| --- | --- | --- | --- | --- |
|  |  |  | 2.5th | 97.5th |
| *virus isolation from blood* |  |  |  |  |
| log peak titre | *μ_V_* | 11.9 | 9.23 | 15.8 |
|  | *s_V_* | 10.3 | 2.90 | 29.5 |
| time of peak titre | *μ_T_* | 1.49 | 1.25 | 1.72 |
|  | *σ_T_* | 0.30 | 0.18 | 0.54 |
| rate for exponential growth phase | 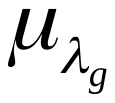 | 6.32 | 4.09 | 12.0 |
|  | 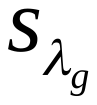 | 3.41 | 1.00 | 9.51 |
| rate for exponential decay phase | 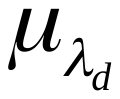 | 5.28 | 4.45 | 6.18 |
|  | 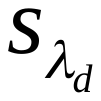 | 86.7 | 13.9 | 374.6 |
| time of clinical onset | *μ_C_* | 1.33 | 1.00 | 1.64 |
|  | *σ_C_* | 0.41 | 0.25 | 0.71 |
| correlation parameter | *ρ_TC_* | 0.96 | 0.58 | 0.99 |
| transmission parameter | *γ* | 0.31 | 0.14 | 0.60 |
| *virus isolation from nasal fluid* |  |  |  |  |
| log peak titre | *μ_V_* | 10.8 | 8.13 | 15.2 |
|  | *s_V_* | 7.92 | 2.21 | 22.7 |
| time of peak titre | *μ_T_* | 1.57 | 1.18 | 1.99 |
|  | *σ_T_* | 0.47 | 0.28 | 0.91 |
| rate for exponential growth phase | 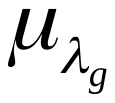 | 9.01 | 4.67 | 25.7 |
|  | 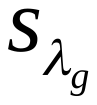 | 1.51 | 0.49 | 3.75 |
| rate for exponential decay phase | 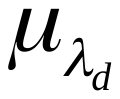 | 3.44 | 2.39 | 5.74 |
|  | 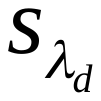 | 6.40 | 1.42 | 26.3 |
| time of clinical onset | *μ_C_* | 1.33 | 1.00 | 1.67 |
|  | *σ_C_* | 0.41 | 0.26 | 0.73 |
| correlation parameter | *ρ_TC_* | 0.91 | 0.24 | 0.99 |
| transmission parameter | *γ* | 0.31 | 0.14 | 0.62 |
| *virus isolation from oesophageal-pharyngeal fluid* | |  |  |  |
| log peak titre | *μ_V_* | 13.0 | 11.4 | 15.2 |
|  | *s_V_* | 57.4 | 12.7 | 254.9 |
| time of peak titre | *μ_T_* | 1.09 | 0.77 | 1.29 |
|  | *σ_T_* | 0.11 | 0.01 | 0.43 |
| rate for exponential growth phase | 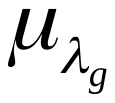 | 5.15 | 4.13 | 7.10 |
|  | 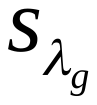 | 105.0 | 11.4 | 409.3 |
| rate for exponential decay phase | 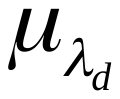 | 2.18 | 1.44 | 3.74 |
|  | 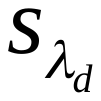 | 6.36 | 1.58 | 28.8 |
| time of clinical onset | *μ_C_* | 1.33 | 0.98 | 1.68 |
|  | *σ_C_* | 0.44 | 0.27 | 0.79 |
| correlation parameter | *ρ_TC_* | -0.11 | -0.90 | 0.87 |
| transmission parameter | *γ* | 0.14 | 0.06 | 0.26 |
