## Supplementary material for "Quantifying the relationship between within-host dynamics and transmission for viral diseases of livestock": S4 Table: Parameter estimates (SwIV)

**S4 Table.** Summary statistics for the marginal posterior densities for hierarchical parameters in a model linking within-host dynamics and transmission of swine influenza virus in pigs.

| parameter | symbol | median | percentiles | |
| --- | --- | --- | --- | --- |
|  |  |  | 2.5th | 97.5th |
| log peak titre | *μ_V_* | 11.6 | 10.2 | 13.2 |
|  | *s_V_* | 33.0 | 11.2 | 87.2 |
| time of peak titre | *μ_T_* | 1.46 | 1.33 | 1.58 |
|  | *σ_T_* | 0.14 | 0.08 | 0.27 |
| rate for exponential growth phase | 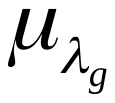 | 4.57 | 3.74 | 5.90 |
|  | 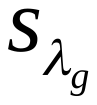 | 49.0 | 6.86 | 322.9 |
| rate for exponential decay phase | 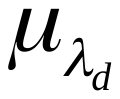 | 3.50 | 1.88 | 4.37 |
|  | 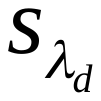 | 35.6 | 7.12 | 230.5 |
| transmission parameter | *γ* | 0.25 | 0.18 | 0.36 |
