## Supplementary material for "Quantifying the relationship between within-host dynamics and transmission for viral diseases of livestock": S5 Table: Prior distributions

**S5 Table.** Prior distributions used when estimating parameters related to the within-host dynamics and transmission of foot-and-mouth disease virus (FMDV) in cattle and swine influenza virus (SwIV) in pigs.

| parameter | symbol(s) | prior | comments |
| --- | --- | --- | --- |
| hierarchical within-host parameters (FMDV and SwIV) | *s_i_*, *μ_i_* (*i*=*V_p_*, *λ_g_*, *λ_d_*), *μ_T_*, *σ_T_* | Exponential(mean=100) | - |
| incubation period parameters (FMDV only) |  |  |  |
|  | *μ_T_* | Gamma(shape=5,mean=1.65) | based on a meta-analysis of challenge experiments [1] |
|  | *σ_T_* | Gamma(shape=3,mean=0.47) |  |
|  | *ρ_TC_* | Uniform(-1,1) | - |
| transmission parameters (FMDV and SwIV) |  |  |  |
| shedding proportional to titre,  transmission parameter common to animals | *γ* | Exponential(mean=10^-4^) | chosen to give *R*_0_ of 10-20  (for FMDV see [2-4]; for SwIV see [5-7]) |
| shedding proportional to log titre,  transmission parameter common to animals | *γ* | Exponential(mean=1) |  |
| shedding proportional to titre,  transmission parameter varies amongst animals | *μ_γ_* | Exponential(mean=-9) |  |
|  | *σ_γ_* | Exponential(mean=1) |  |
| shedding proportional to log titre,  transmission parameter varies amongst animals | *s_γ_* | Exponential(mean=1) |  |
|  | *μ_γ_* | Exponential(mean=1) |  |
| error standard deviation (FMDV and SwIV) | *σ_E_* | Gamma(shape=5,mean=0.3) | - |
