## Supplementary material for "Quantifying the relationship between within-host dynamics and transmission for viral diseases of livestock": S6 Table: Approximation accuracy

**S5 Table.** Accuracy of heuristic approximations for the individual reproduction number and generation time.

| virus and viral titre | median (99% range) for exact minus approximate expression | |
| --- | --- | --- |
|  | reproduction number | generation time |
| FMDV, blood | -0.016 (-0.049, 0.032) | 0.001 (-0.033, 0.003) |
| FMDV, nasal fluid | -0.008 (-0.167, 0.148) | 0.003 (-0.081, 0.031) |
| FMDV, OP fluid | -0.007 (-0.024, 0.023) | 0.005 (-0.0004, 0.015) |
| SwIV, nasal swabs | -0.021 (-0.037, -0.008) | 0.002 (-0.003, 0.008) |
