## Supplementary figures and images for "Quantifying the relationship between within-host dynamics and transmission for viral diseases of livestock"

### S1 Figure

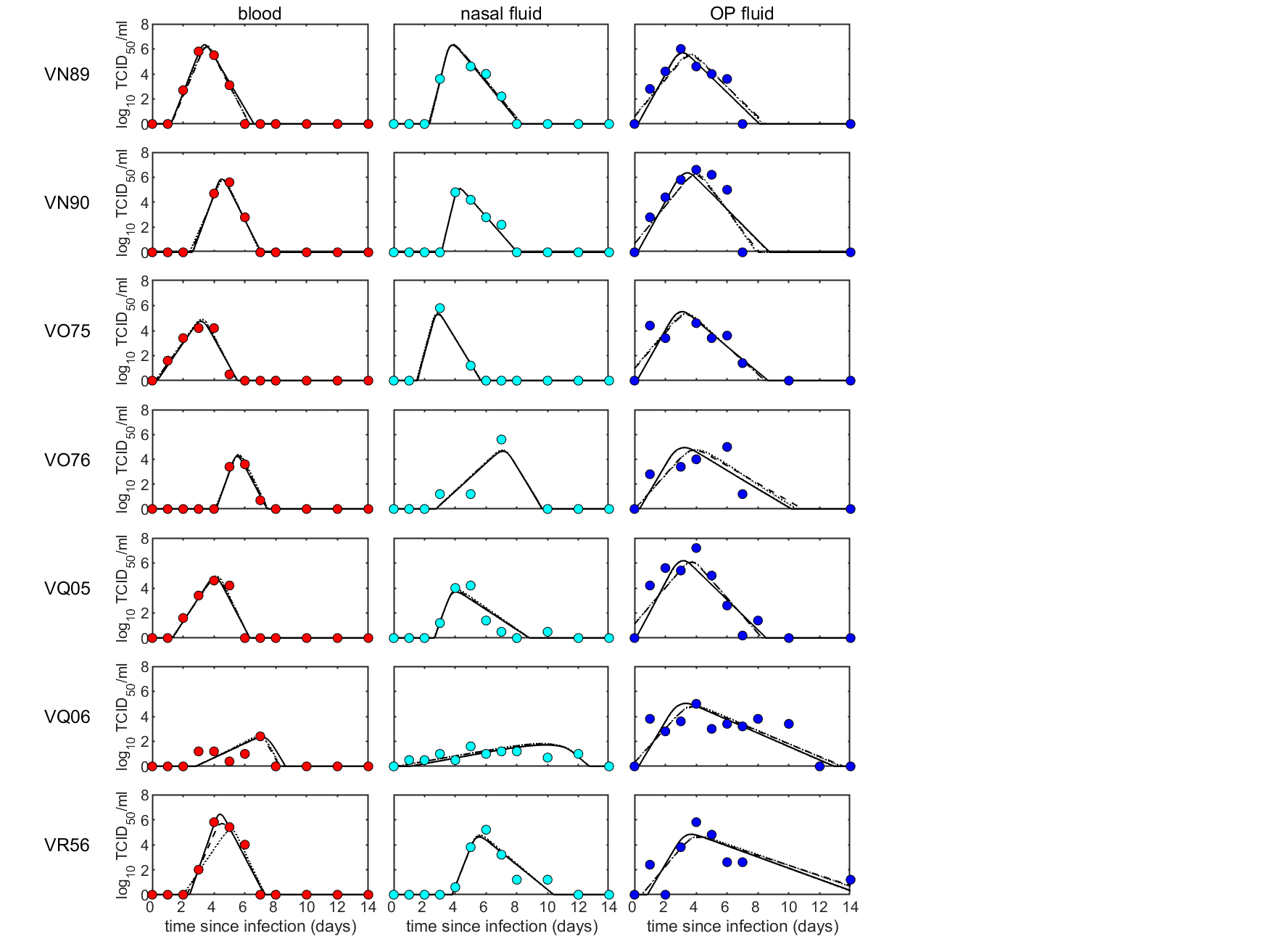

### S2 Figure

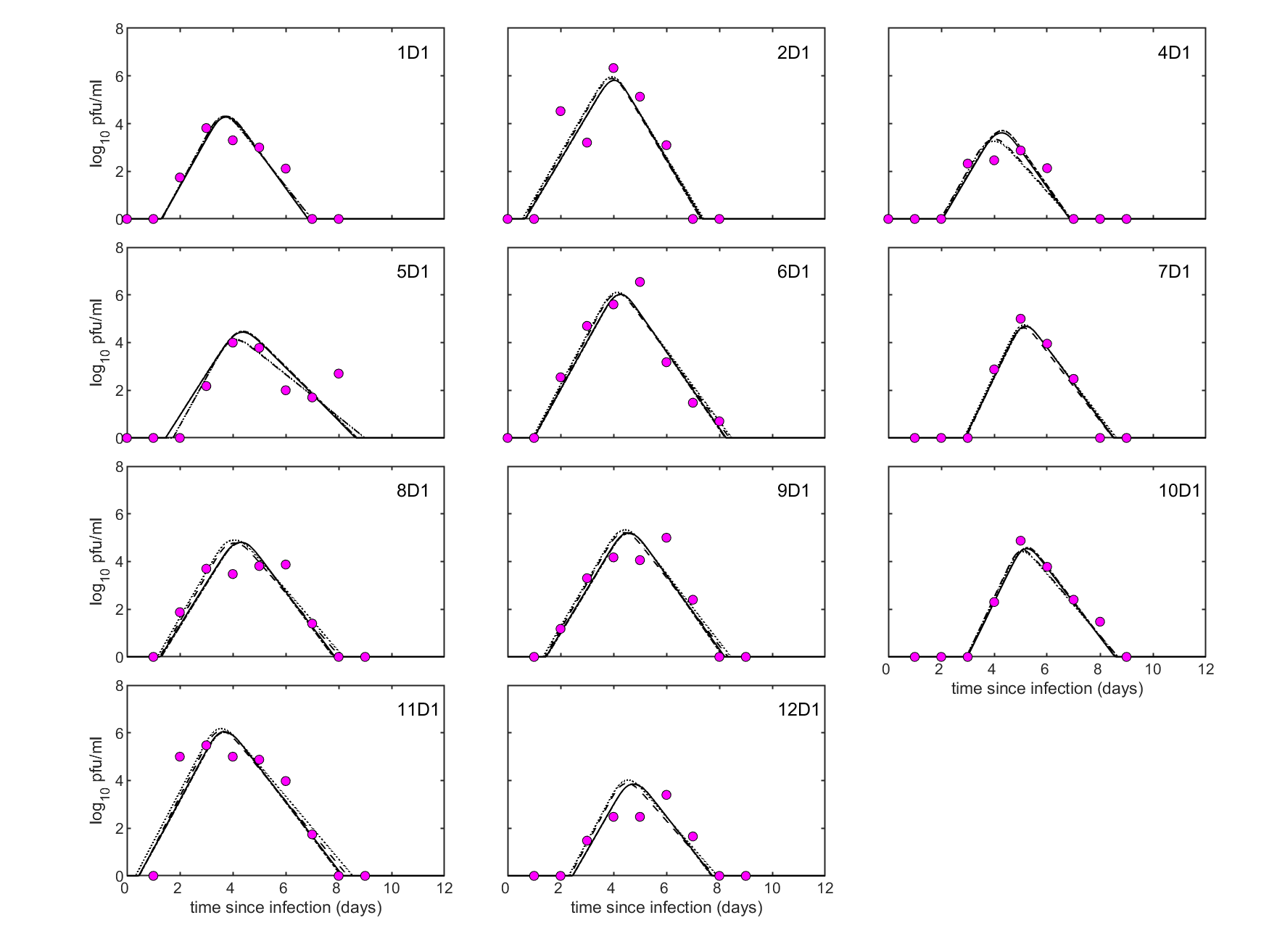

### S3 Figure

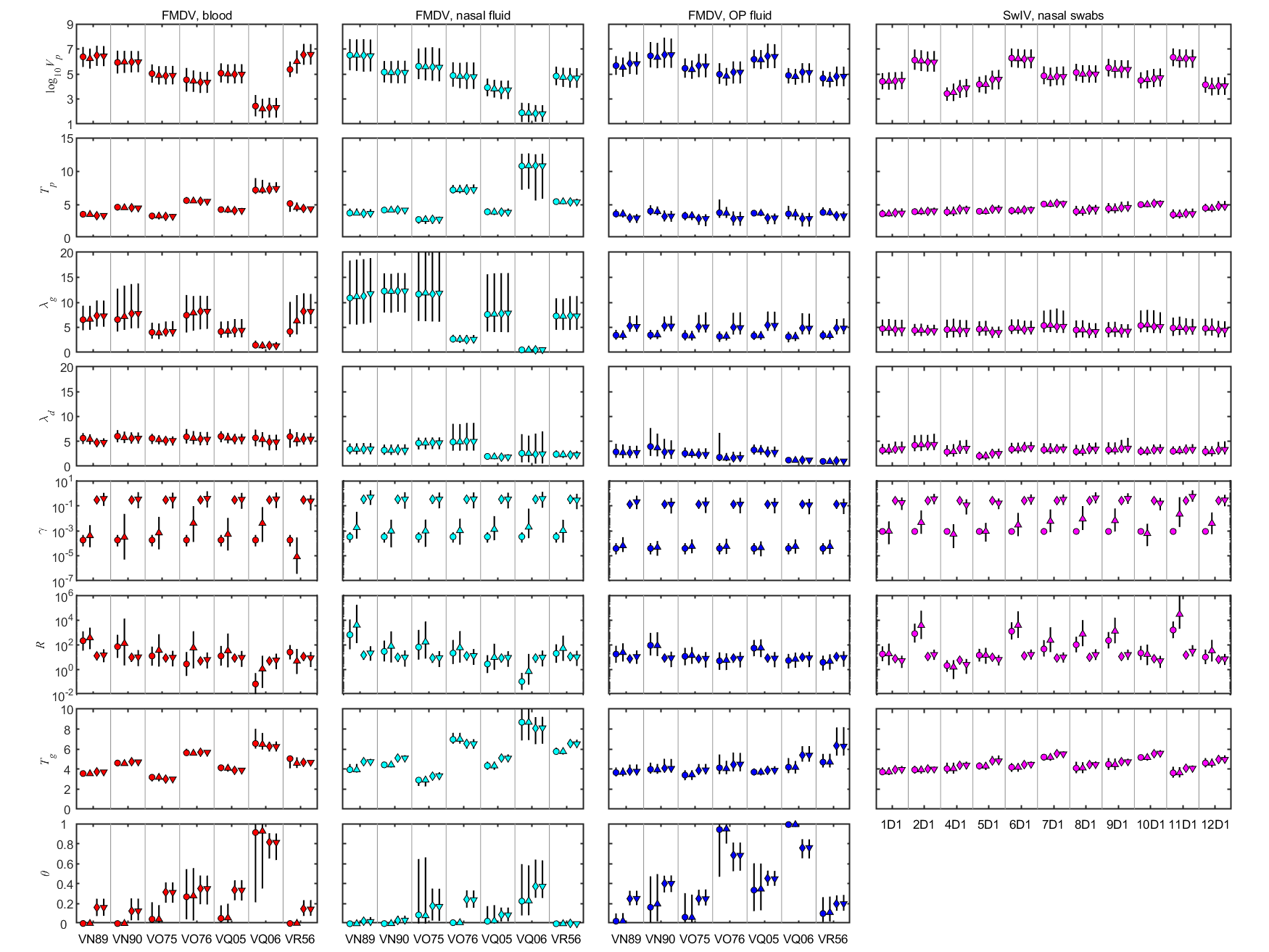

### S4 Figure

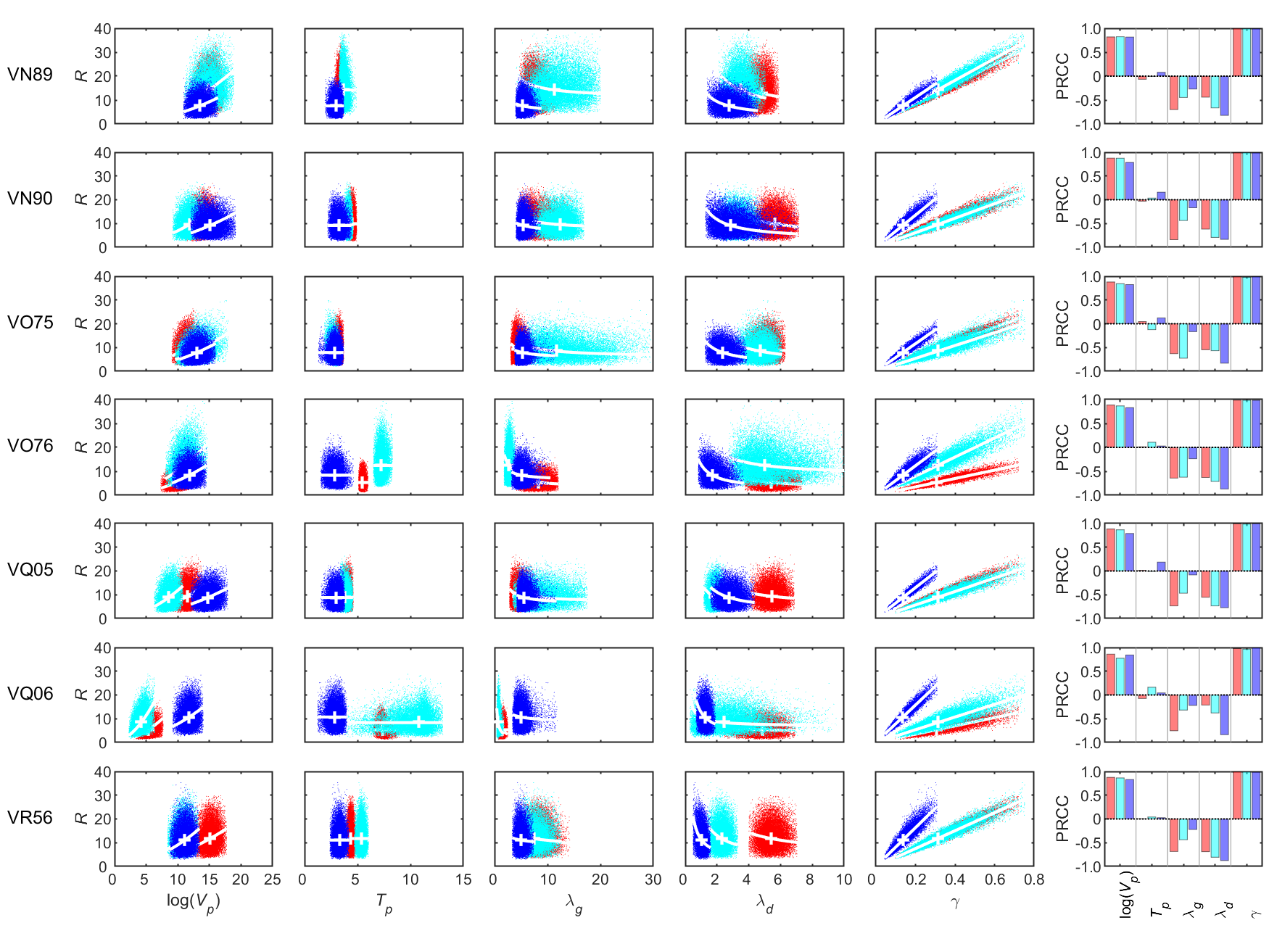

### S5 Figure

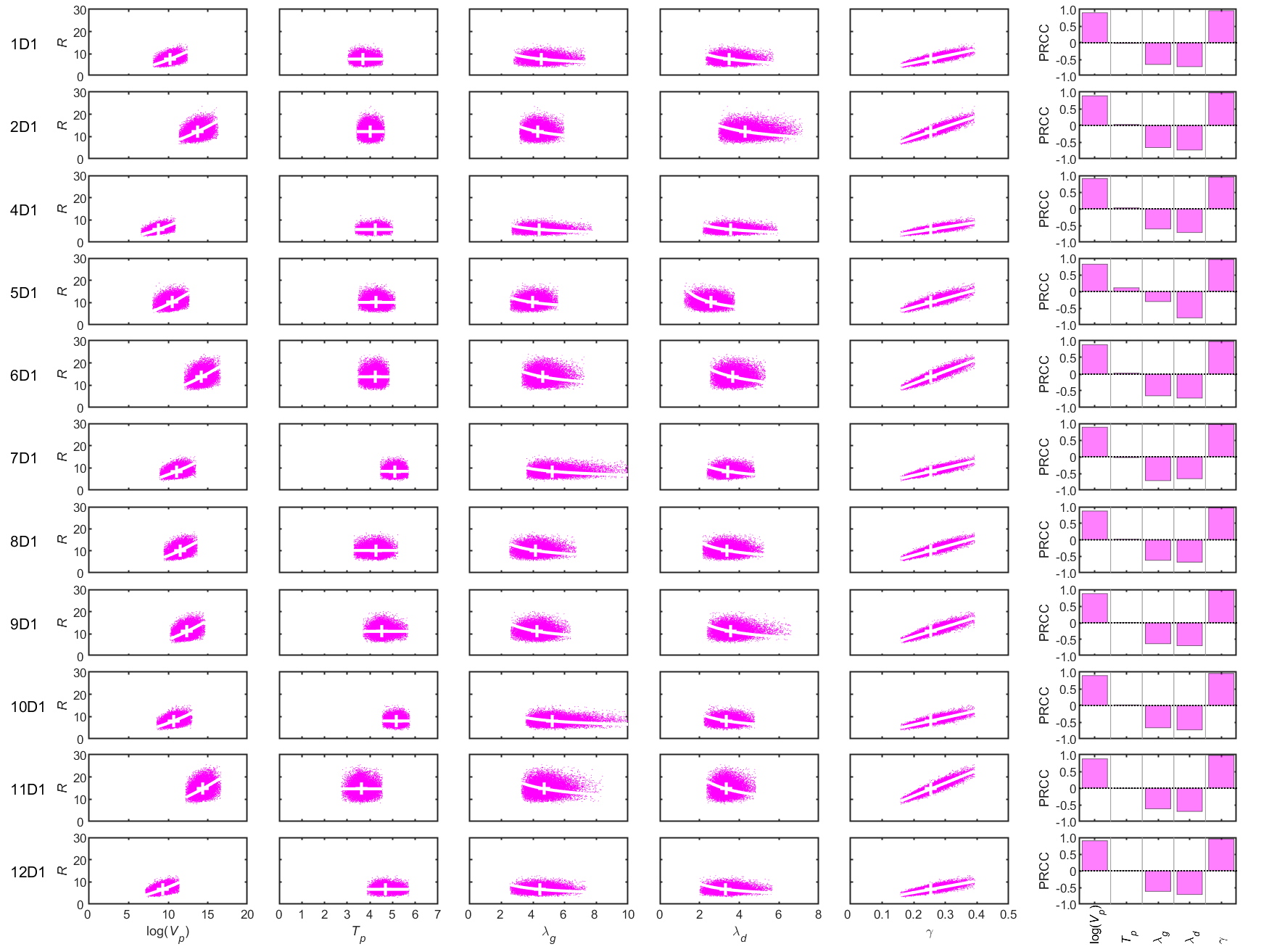

### S6 Figure

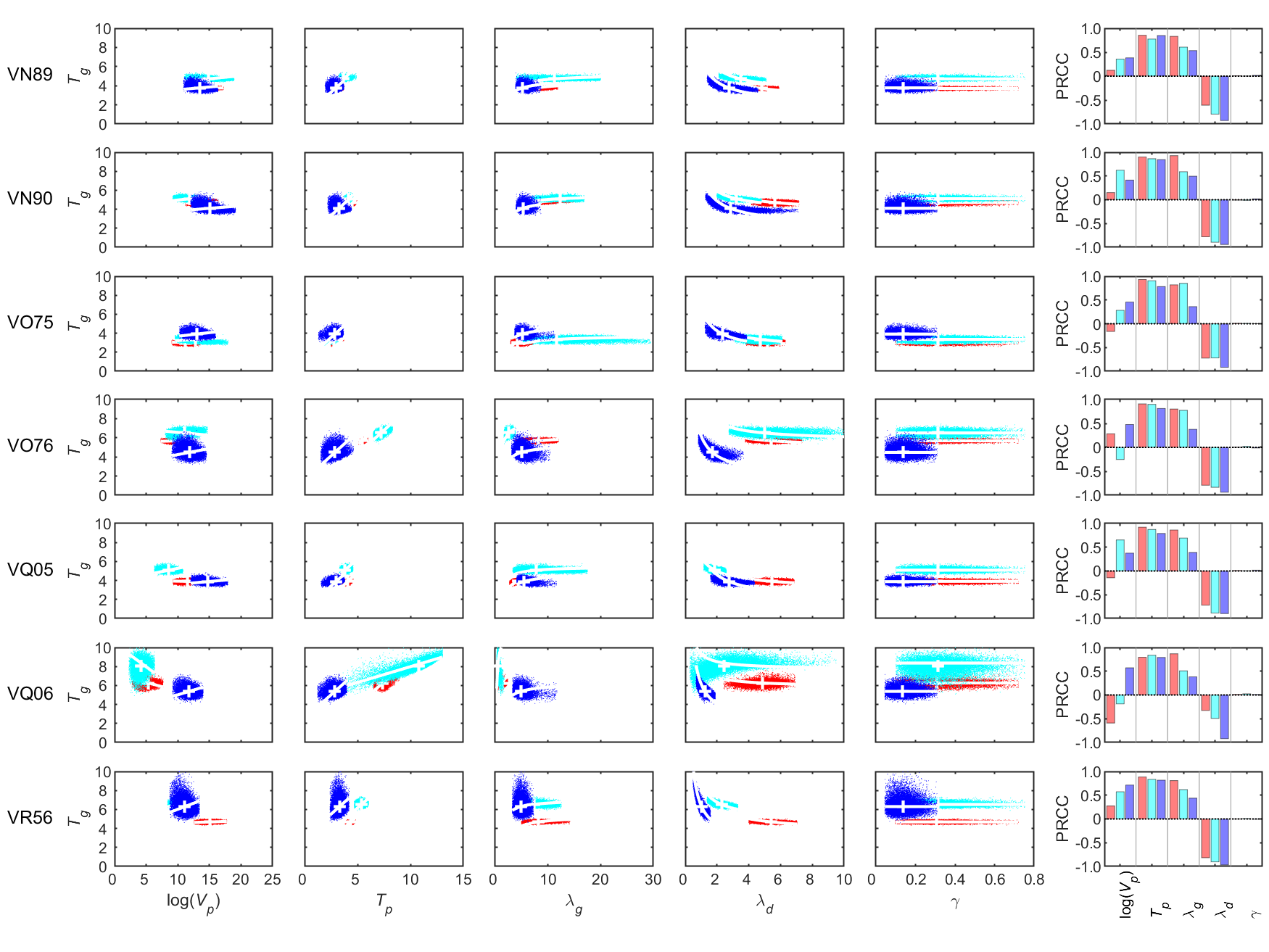

### S7 Figure

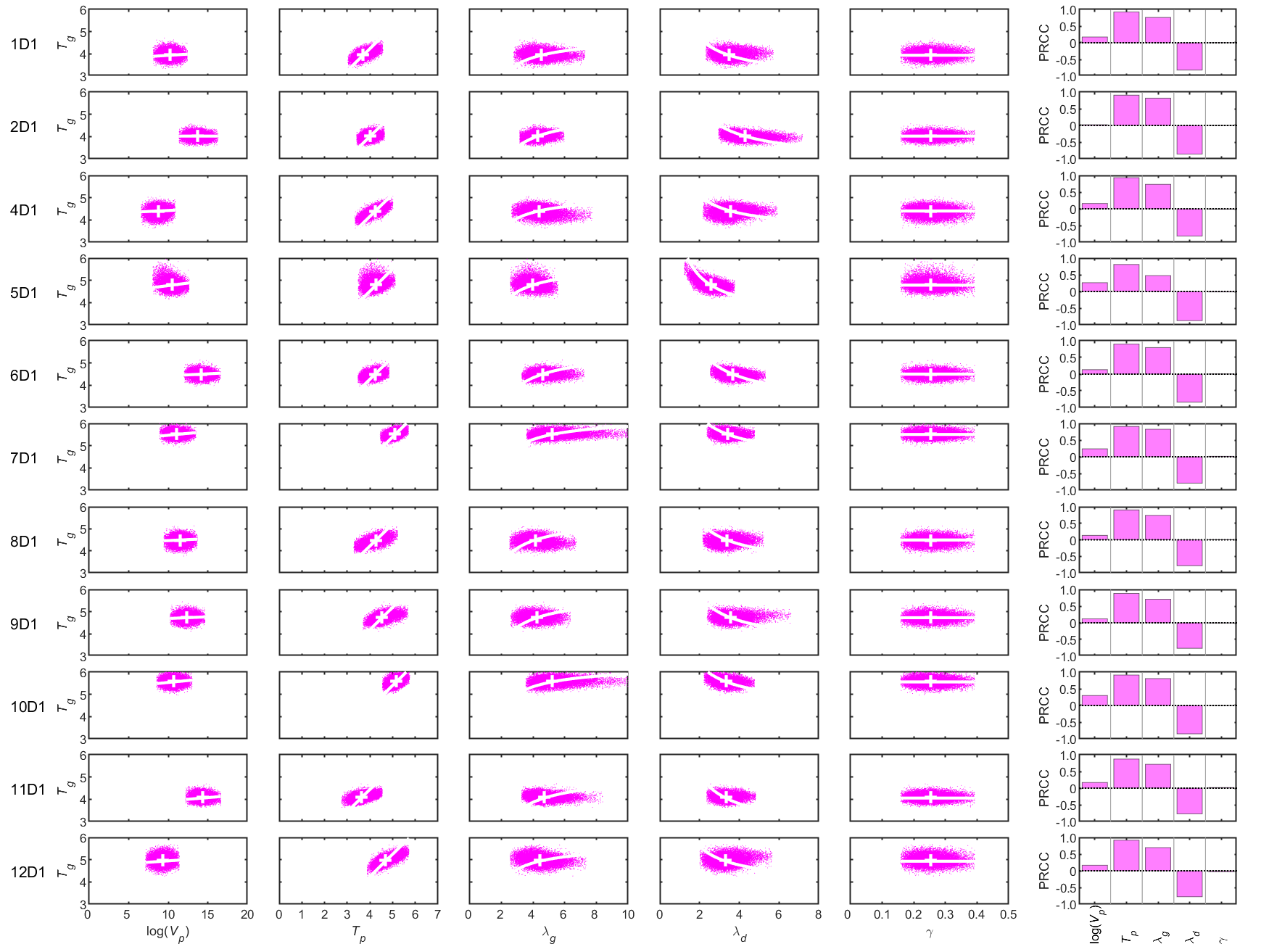

### S8 Figure

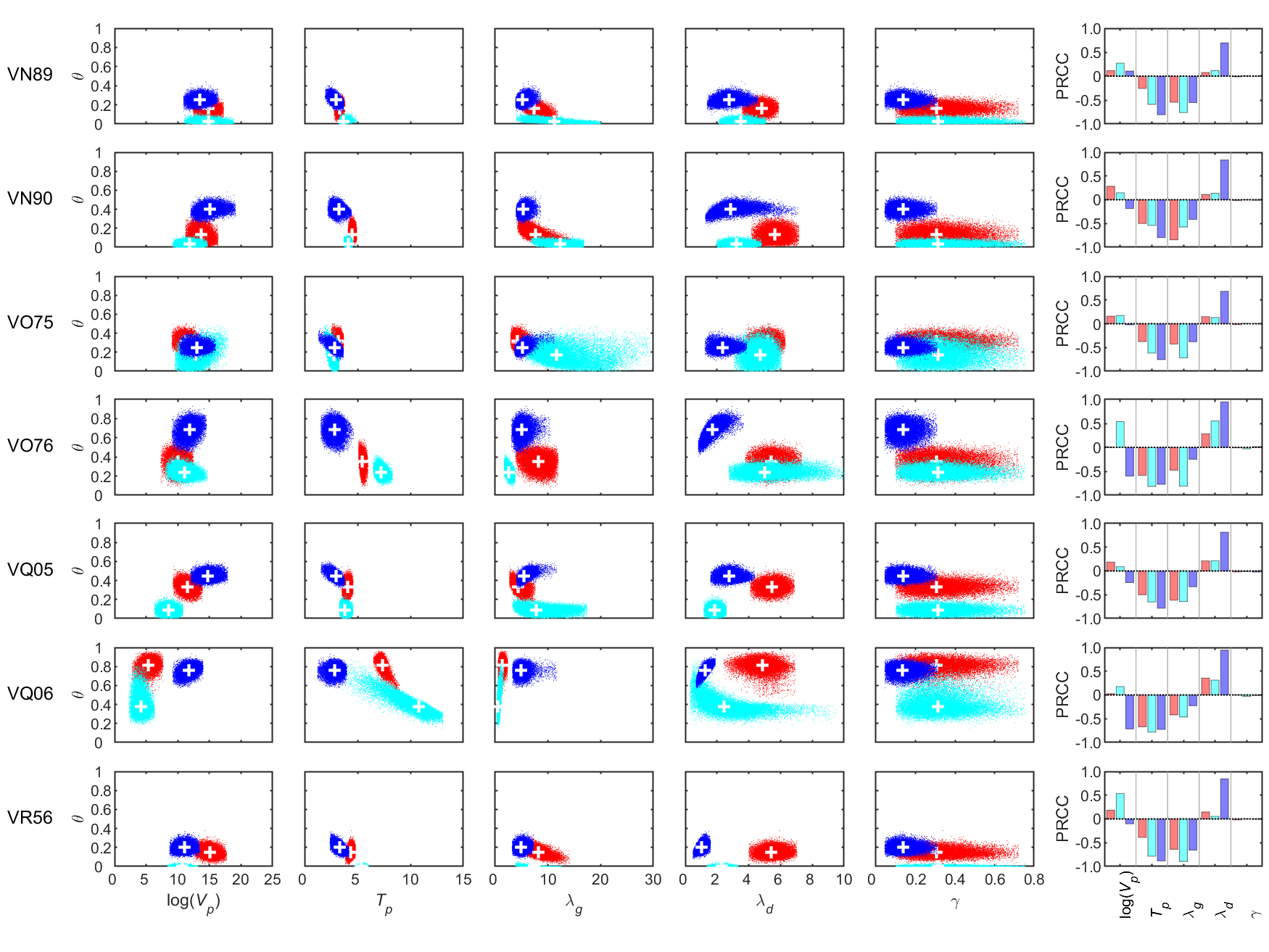

### S9 Figure

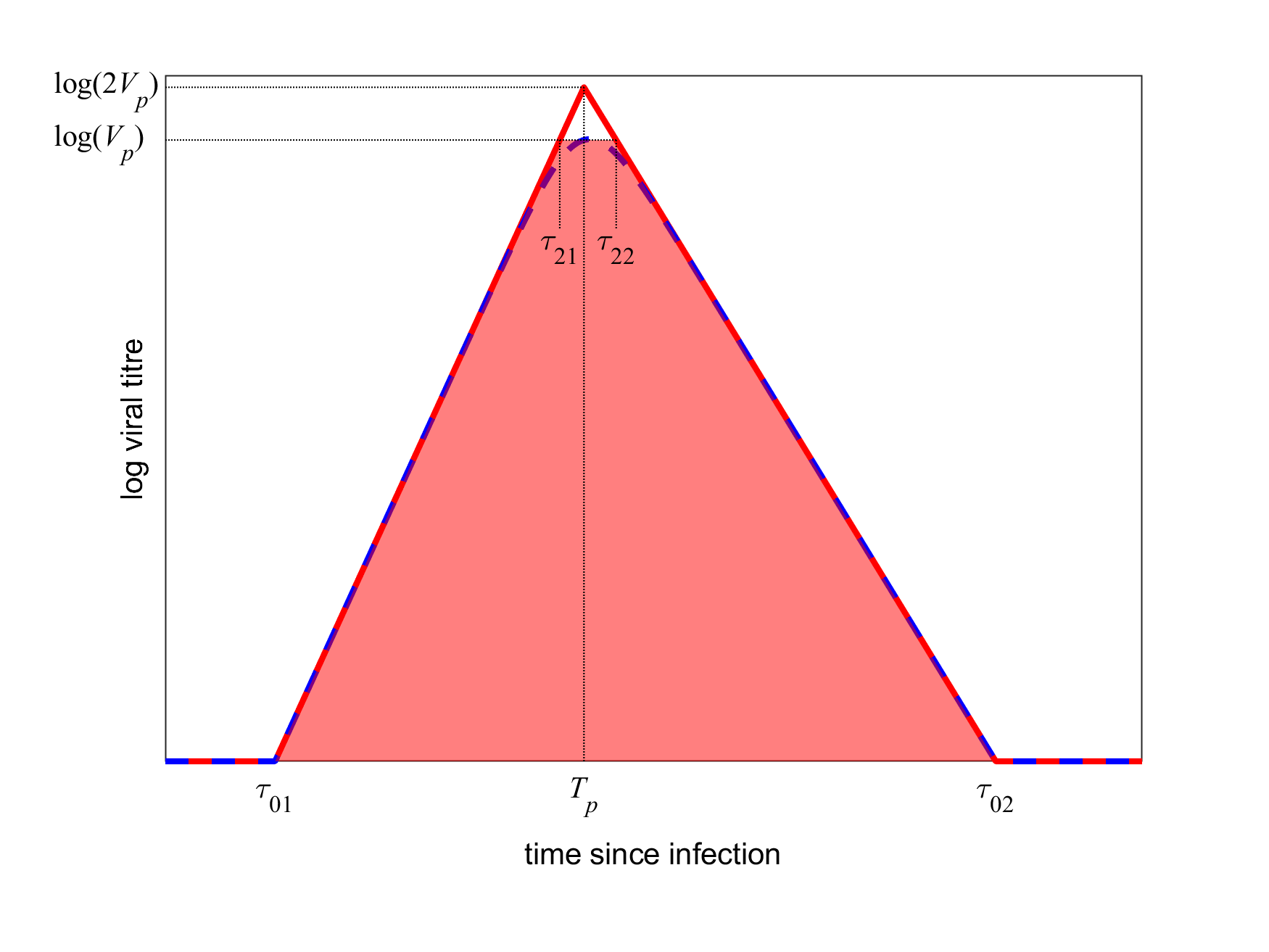
